# Fast and sensitive taxonomic assignment to metagenomic contigs

**DOI:** 10.1101/2020.11.27.401018

**Authors:** M. Mirdita, M. Steinegger, F. Breitwieser, J. Söding, E. Levy Karin

## Abstract

MMseqs2 taxonomy is a new tool to assign taxonomic labels to metagenomic contigs. It extracts all possible protein fragments from each contig, quickly retains those that can contribute to taxonomic annotation, assigns them with robust labels and determines the contig’s taxonomic identity by weighted voting. Its fragment extraction step is suitable for the analysis of all domains of life. MMseqs2 taxonomy is 2-18x faster than state-of-the-art tools and also contains new modules for creating and manipulating taxonomic reference databases as well as reporting and visualizing taxonomic assignments.

**Availability:** MMseqs2 taxonomy is part of the MMseqs2 free open-source software package available for Linux, macOS and Windows at https://mmseqs.com.

## I. INTRODUCTION

Metagenomic studies shine a light on previously unstudied parts of the tree of life. However, unraveling taxonomic composition accurately and quickly remains a challenge. While most methods label short metagenomic reads (reviewed in [11]), only a handful (e.g. [6]) assign entire contigs, even though this should lead to improved accuracy.

Recently, [12] developed CAT, a tool for taxonomic annotation of contigs based on protein homologies to a reference database. It combines Prodigal [7] for predicting open reading frames (ORFs), DIAMOND [3] to search with the translated ORFs, and logic to aggregate individual ORF annotations. CAT achieved higher precision than state-of-the-art tools on bacterial benchmarks. Despite its advantage over existing methods, CAT has limitations: (1) Prodigal was designed for prokaryotes and not eukaryotes [13]; (2) Prodigal runs single-threaded, limiting applicability to metagenomics; (3) CAT’s *r* parameter determines the cut-off score below each ORF’s top-hit above which hits are included in the ORF’s lowest common ancestor (LCA) computation. Although the authors provide guidelines to set *r*, it is unclear how general they are.

Here we present MMseqs2 taxonomy, a novel protein-search-based tool for taxonomy assignment to contigs. It overcomes the aforementioned limitations by extracting all possible protein fragments, covering the coding repertoire of all domains of life. It quickly eliminates fragments that do not bear minimal similarity to the reference database, and searches with the remaining ones. MM-seqs2 taxonomy uses an approximate 2bLCA [5] strategy to assign translated fragments to taxonomic nodes (Supp. Inf.). The hits for the a2bLCA computation are determined automatically, saving the need to tune an equivalent of CAT’s *r* parameter. It outperforms CAT on bacterial and eukaryotic data sets.

## II. METHODS

### Input

Contigs are provided as (compressed) FASTA/Q files. As reference, the *databases* workflow can download and prepare various public taxonomy databases, such as, nr [1], UniProt [2] or GTDB [10]. Alternatively, users can prepare their own taxonomic reference database (see MMseqs2 wiki).

### Algorithm

The four main steps are described in Fig. 1A.

**FIG. 1.**
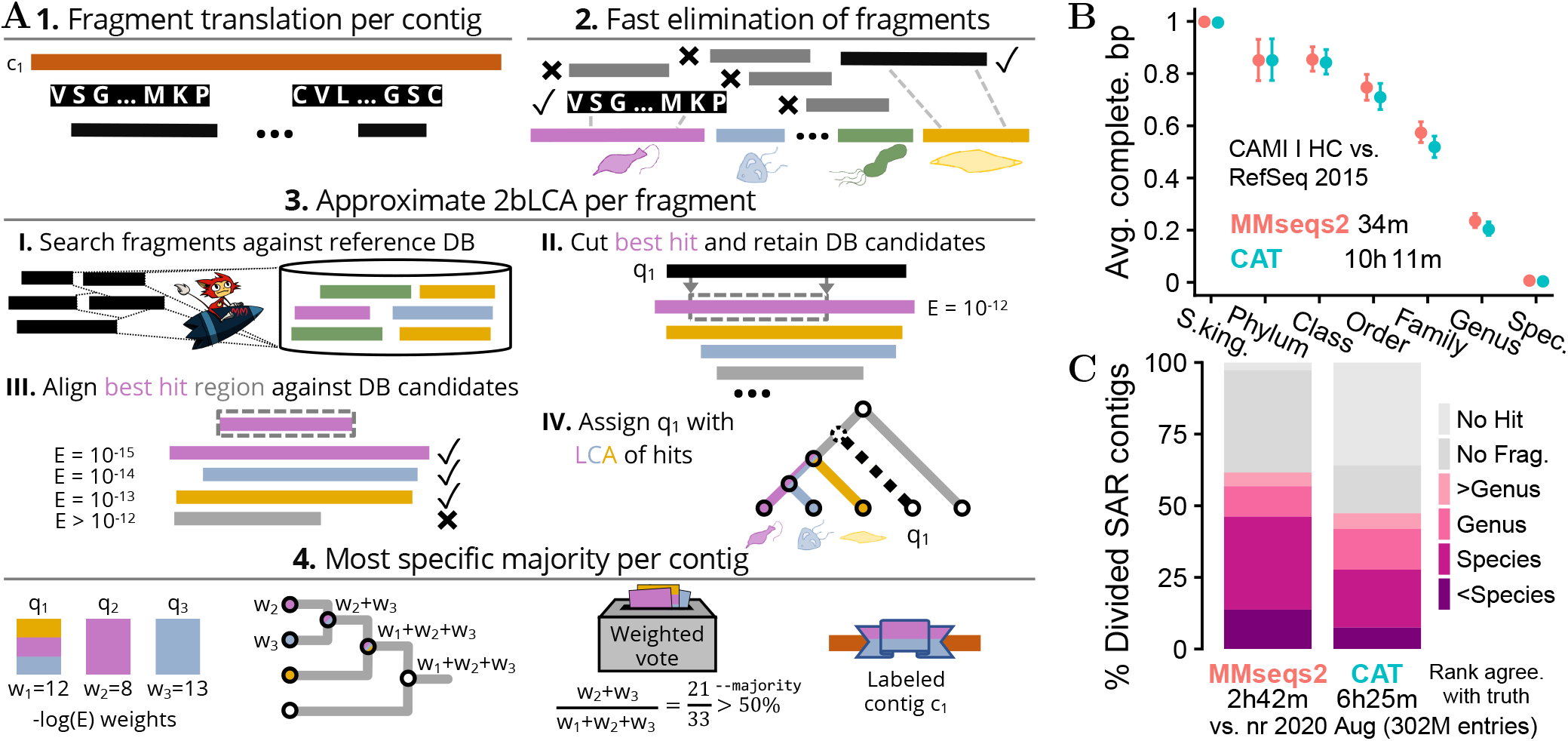
(**A**) Taxonomy assignment algorithm in four steps: (1) Translate all possible protein fragments in six frames from all contigs. (2) Reject fragments unlikely to find a taxonomic hit in later stages (full details in Supp. Inf.). (3) Assign taxonomic nodes using an approximate 2bLCA procedure. Each query fragment *q* is searched against the reference database, resulting in a list *l* of all its homologous targets. The aligned region between *q* and the best hit *t* (with *E*-value *E*(*q, t*)) is aligned against all targets in *l*. We assign *q* the LCA of the taxonomic lables of all target sequences that have an *E*-values lower than *E*(*q, t*). By realigning *l* we could avoid the costly second search of 2bLCA. (4) Each assigned *q* contributes its weight (–log *E*(*q, t*)) to its taxonomic label and all labels above it, up to the root. The contig’s taxonomic node is determined as the most specific taxonomic label, which has a support of at least the `--majority` parameter. The support of a label is the sum of its contributing weights divided by the total sum of weights. (**B**) MMseqs2 taxonomy (red) is ~18x faster and achieves similar average completeness to CAT (turquoise) on a bacterial benchmark. (**C**) MMseqs2 assigns taxons to eukaryotic SAR contigs more accurately than CAT across all phylogenetic levels, at twice the speed. At species level, MMseqs2 taxonomy classifies 46% contigs correctly versus 28% for CAT. Runtimes measured on a 2×14-core Intel E5-2680v4 server with 768GB RAM.

### Output

MMseqs2 taxonomy returns the following eight fields for each contig accession: (1) the taxonomic identifier (taxid) of the assigned label, (2) rank, (3) name, followed by the number of fragments: (4) retained, (5) taxonomically assigned, and (6) in agreement with the contig label (i.e., same taxid or have it as an ancestor), (7) the support the taxid received and, optionally, (8) the full lineage. The result can be converted to a TSV-file, and to a Kraken [14] report or a Krona [9] visualization (Supp. Information).

## III. RESULTS

### Bacterial dataset

The CAMI-I high-complexity challenge and its accompanying RefSeq 2015 reference database [11] were given to MMseqs2 and CAT. AM-BER v2 [8] was used to assess the taxonomic assignment by computing the average completeness (Fig 1B) and purity (Fig S1) bp using its taxonomic binning benchmark mode. At similar assignment quality, MMseqs2 taxonomy is 18x faster than CAT. Using the nr database, MM-seqs2 is 10x faster (Fig S2).

### Eukaryotic dataset

All 57 SAR (taxid 2698737) RefSeq assemblies and their taxonomic labels were downloaded from NCBI in 08/2020. To resemble metagenomic data, their scaffolds were randomly divided following the length distribution of contigs assembled for sample ERR873969 of eukaryotic Tara Oceans [4], resulting in 2.7 million non-overlapping contigs with a minimal length of 300 bp. Using nr from 08/2020, MMseqs2 classified more contigs than CAT (62% vs. 47%). For 36%, CAT extracted a fragment that did not hit the reference, suggesting fragments extracted by MMseqs2 are more informative for eukaryotic taxonomic annotation (Fig 1C, S3).

## IV. CONCLUSION

MMseqs2 taxonomy is as accurate as CAT on a bacterial data set while being 3-18x faster and requiring fewer parameters. Its extracted fragments make it suitable for analyzing eukaryotes. It is accompanied by several taxonomy utility modules to assist with taxonomic analyses.

## FUNDING

ELK is a FEBS long-term fellowship recipient. The work was supported by the BMBF CompLifeSci project horizontal4meta; the ERC’s Horizon 2020 Framework Programme [‘Virus-X’, project no. 685778]; the National Research Foundation of Korea grant funded by the Korean government (MEST) [2019R1A6A1A10073437, NRF-2020M3A9G7103933]; and the Creative-Pioneering Researchers Program through Seoul National University.

### Conflict of Interest

none declared

## Notes

### Competing Interest Statement

The authors have declared no competing interest.

## REFERENCES

[1] Agarwala, R. et al (2018). Database resources of the National Center for Biotechnology Information. Nucleic Acids Res., 46(D1), D8–D13.

[2] Bateman, A. (2019). UniProt: a worldwide hub of protein knowledge. Nucleic Acids Res., 47(D1), D506–D515.

[3] Buchfink, B. et al (2015). Fast and sensitive protein alignment using DIAMOND. Nat. Methods, 12(1), 59–60.

[4] Carradec, Q. et al (2018). A global ocean atlas of eukaryotic genes. Nat. Commun., 9(1), 373.

[5] Hingamp, P. et al (2013). Exploring nucleo-cytoplasmic large DNA viruses in Tara Oceans microbial metagenomes. ISME J., 7(9), 1678–1695.

[6] Huson, D.H. et al (2018). MEGAN-LR: new algorithms allow accurate binning and easy interactive exploration of metagenomic long reads and contigs. Biol. Direct, 13(1), 6.

[7] Hyatt, D. et al (2010). Prodigal: prokaryotic gene recognition and translation initiation site identification. BMC Bioinform., 11(1), 119.

[8] Meyer, F. et al (2018). AMBER: Assessment of Metagenome BinnERs. Gigascience, 7(6), giy069.

[9] Ondov, B.D. et al (2011). Interactive metagenomic visualization in a Web browser. BMC Bioinform., 12(1), 385.

[10] Parks, D.H. et al (2020). A complete domain-to-species taxonomy for Bacteria and Archaea. Nat. Biotechnol., 38(9), 1079–1086.

[11] Sczyrba, A. et al (2017). Critical Assessment of Metagenome Interpretation—a benchmark of metagenomics software. Nat. Methods, 14(11), 1063–1071.

[12] von Meijenfeldt, F.A.B. et al (2019). Robust taxonomic classification of uncharted microbial sequences and bins with CAT and BAT. Genome Biol., 20(1), 217.

[13] West, P.T. et al (2018). Genome-reconstruction for eukaryotes from complex natural microbial communities. Genome Res., 28(4), 569–580.

[14] Wood, D.E. et al (2019). Improved metagenomic analysis with Kraken 2. Genome Biol., 20(1), 257.

