## Supplement for "Fast and sensitive taxonomic assignment to metagenomic contigs"

### Supplementary Material for Fast and sensitive taxonomic assignment to metagenomic contigs using MMseqs2


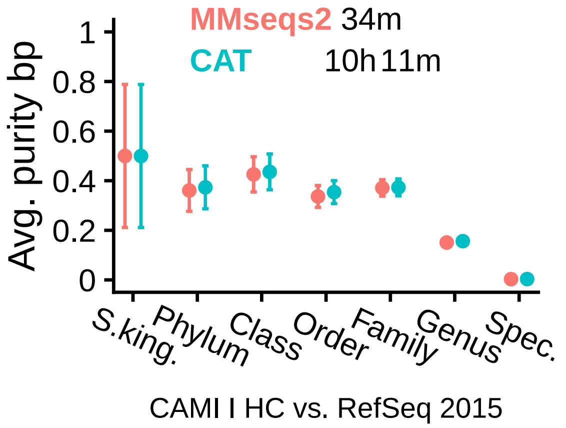


FIG. S1. MMseqs2 (red) is ~18x faster and achieves similar average purity bp to CAT (turquoise) on a bacterial benchmark. The CAMI taxonomic reference database does not contain any species, which were included in the challenge. All classifiers score 0% bp completeness/purity at that rank. Runtimes measured on a server with 2x14-core Intel E5-2680v4 CPUs and 768GB RAM.

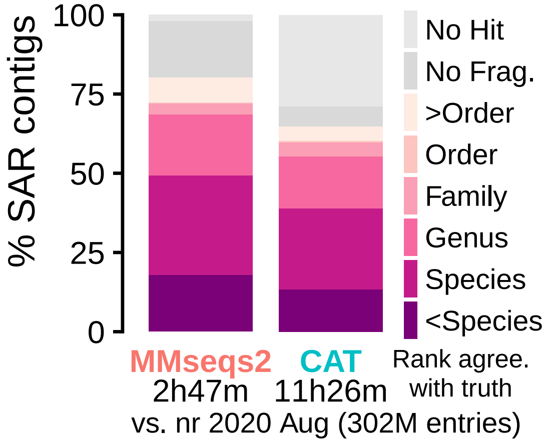


FIG. S3. The fragments extracted and retained by MMseqs2 from 66,630 eukaryotic scaffolds result in more correctly classified scaffolds than those extracted for CAT by Prodigal. All runtimes were measured on a server with two 14-core Intel E5-2680v4 CPUs and 768GB RAM.

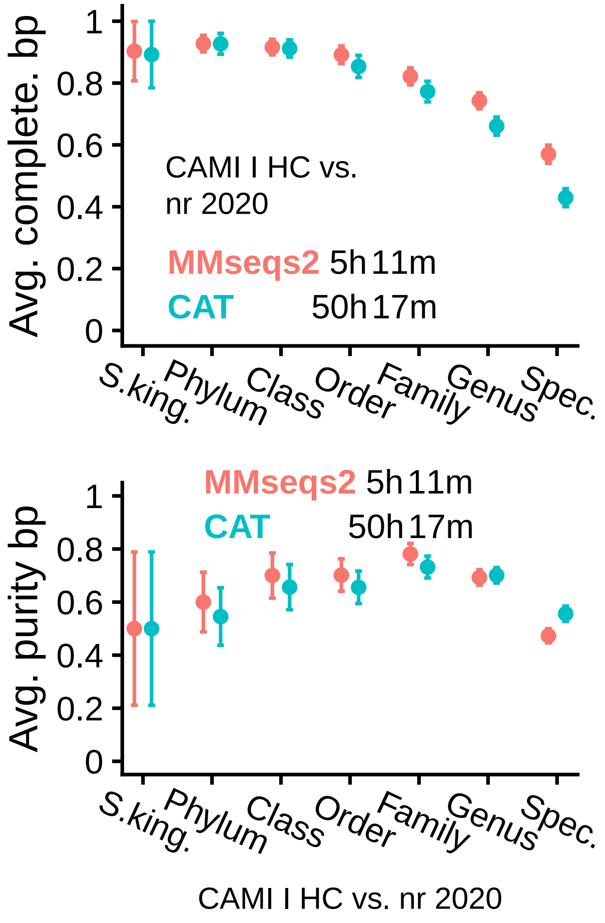


FIG. S2. MMseqs2 (red) is ~10x faster and achieves similar accuracy to CAT (turquoise) on a bacterial benchmark, using the nr as reference (16x more entries than RefSeq 2015). In contrast to Fig 1B and Fig S1, the nr database does not exclude any species, which allows for correct classifications at the species level. All runtimes were measured on a server with two 14-core Intel E5-2680v4 CPUs and 768GB RAM.

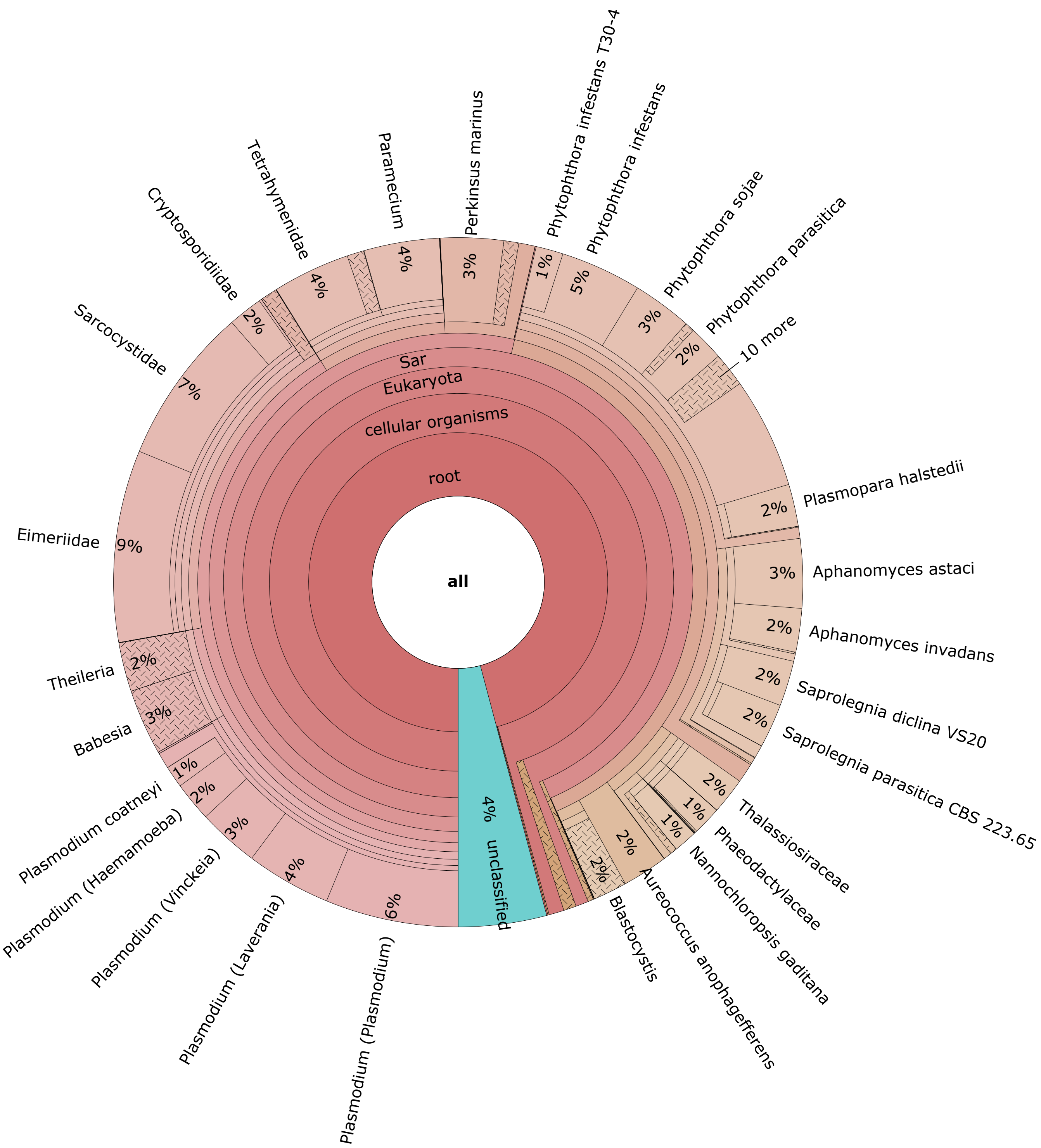


FIG. S4. Krona visualization of the classified contigs of Fig. 1C (62% of all contigs) generated using `mmseqs taxonomyreport --report-mode 1`.

#### EARLY PROTEIN FRAGMENT REJECTION

MMseqs2 taxonomy uses the `prefilter` module to find the translated fragment-to-reference match with the highest number of consecutive similar $k$-mer matches on the same diagonal. Fragments with fewer than three matches to any reference sequence are removed. The remaining pairs are aligned without gaps using `rescorediagonal` and the fragment in each pair is retained if the pair's E-value is smaller than 100.

#### APPROXIMATE 2BLCA

The 2bLCA procedure consists of two searches: (I) A search with a query sequence against a set of target sequences. (II) A search with the aligned region of the most significant sequence match against the same target sequences. The taxonomic labels of all hits with an E-value smaller or equal to the best hit E-value in search I are used to compute an LCA.

The prefilter of MMseqs2 can quickly identify candidates of homology, which are then verified by a costly alignment step. We approximate 2bLCA by assuming that most candidates found in a prefiltering step of search I would also cover the candidates found by search II. Thus, we reuse the same list of prefiltering candidates for both alignment steps.

Additionally, we exploit MMseqs2's support for multiple alignment modes to calculate only the score and E-value, or to additionally compute the alignment boundaries. In the initial alignment of the query against the target candidates, we use the first mode to find the hit with the best E-value and then we recompute, for the best hit only, the alignment boundaries with the slower, second alignment mode. To compute the E-values of the best matching aligned region to the target sequences we also use the first, faster alignment mode.

We applied `--max-accept 30` and `--max-rejected 5` in the first alignment and `--max-rejected 5` in the second alignment, to further speed up the alignments.

| Name | Version | Comment |
|---|---|---|
| MMseqs2 | Git: 7da33b0 | Benchmarks |
| MMseqs2 | Git: 6379422 | nr DB creation |
| CAT | v5.1.1 | |

TABLE S1. Software versions used in this manuscript.

| Sequence set | Version | Entries | Residues/Bases |
|---|---|---|---|
| CAMI I HC GSA | 2015 | 42k | 2.8B nucl. |
| CAMI I RefSeq | 2015 | 16M | 6.5B aa. |
| SAR (unchopped) | 08/2020 | 67k | 2.2B nucl. |
| SAR (chopped) | 08/2020 | 2.7M | 2.2B nucl. |
| nr | 08/2020 | 303M | 109B aa. |

TABLE S2. Sequence sets used in this manuscript.

| Method | Target DB | Peak RAM |
|---|---|---|
| MMseqs2 | CAMI I RefSeq | 60 GB |
| MMseqs2 | nr | 253 GB |
| CAT | CAMI I RefSeq | 45 GB |
| CAT | nr | 63 GB |

TABLE S3. Peak RAM use of MMseqs2 and CAT with CAMI I HC dataset. Memory use was measured on a server with two 14-core Intel E5-2680v4 CPUs and 768GB RAM. Note, however, that both methods can split the database into chunks and search them one after the other to adapt to available system memory.
